# Decoding the anti-aging effect of retinol in reshaping the human skin microbiome niches

**DOI:** 10.1101/2024.06.26.600860

**Authors:** Minyan Gui, Jingmin Cheng, Xueni Lin, Danni Guo, Qi Zhou, Wentao Ma, Hang Yang, Xueqing Chen, Zhao Liu, Lan Ma, Xinhui Xing, Peng Shu, Xiao Liu

**Affiliations:** The Institute of Biopharmaceutical and Health Engineering, Tsinghua Shenzhen International Graduate School, Tsinghua University, Shenzhen, Guangdong, China; HBN Research Institute and Biological Laboratory, Shenzhen Hujia Technology Co., Ltd., 518000, Shenzhen, Guangdong, China; State Key Laboratory Basis of Xinjiang Indigenous Medicinal Plants Resource Utilization, Xinjiang Technical Institute of Physics and Chemistry, Chinese Academy of Sciences, 830011, Urumqi, Xinjiang, China; University of Chinese Academy of Sciences, 100049, Beijing, China

**Author notes:** these authors contributed equally in this work (Minyan Gui, Jingmin Cheng, Xueni Lin, Danni Guo). Correspondence goes to Xiao Liu and Peng Shu.

## Abstract

Retinol has been widely added to skincare products due to its ability to promote the proliferation of skin keratinocytes and regulate skin cell collagen expression. While it is known the skin harbors a myriad of commensal bacteria, the impact of retinol on the skin microbiome, as well as the role of the skin microbiome in mediating the anti-aging properties of retinol, remains poorly understood. In this study, we incorporated phenomics, metagenomics and metabolomics to explore the human skin alterations during the anti-aging process mediated by retinol, and potential interactions between retinol, skin microbiome and metabolites.

Topical retinol significantly improved skin conditions, including enhancing skin hydration, acidifying the epidermis, strengthening the skin barrier, and reducing the number and volume of wrinkles. Furthermore, retinol also reshaped the skin microecology by altering the structure and function of the skin microbiome as well as the host and microbial metabolites. Through GEM construction, we identified 2 skin microorganism, *Sericytochromatia sp.* and *Corynebacterium kefirresidentii* capable of oxidizing retinol to retinal. Over 10 skin microbes can utilize UDP-glucose as a carbon source, potentially accelerating RAG hydrolysis and increasing glucuronic acid consumption. The retinoic acid and retinol generated by RAG hydrolysis are reused by skin cells and microbes, enhancing retinol metabolism and its effective duration. This combined effect between the skin microbiome and retinol improves skin condition and anti-aging efficacy.

## Introduction

Retinol is a derivative of vitamin A and a precursor of retinoic acid but with a milder effect and better tolerated by the skin^1^. Unlike retinoic acid, which is a strong skin irritant^2,3^, retinol is more suitable for sensitive skin and is less likely to cause adverse reactions such as dryness, erythema, and itching^4,5^. Therefore, retinol is widely added to cosmetics and skin care products as an anti-aging active ingredient^4,6–8^. Being a fat-soluble substance, retinol can penetrate skin cells and be metabolized to retinoic acid, the bioactive form of retinoids^9^, in a two-step reaction^10,11^. Firstly, alcohol dehydrogenases (ADHs) convert retinol to retinaldehyde, which can be subsequently oxidized to retinoic acid by retinaldehyde dehydrogenases (RALDH1, RALDH2, RALDH3)^9,10^. Retinoic acid can bind to retinoic acid receptors^1,12–14^ and retinoid X receptors^15–17^ to achieve regulatory functions.

Retinol has demonstrated its effectiveness in promoting wrinkle conditions in human skin. The anti-aging mechanism of retinol has been widely studied^12,17–19^. It is reported that retinol can induce epidermal thickening and stratum corneum renewal by activating IFE stem cells and promoting the proliferation and differentiation of keratinocytes^20–22^. It can also increase the expression of collagen, fibronectin, and tropoelastin^18^, which are the main structural proteins in skin, in dermal fibroblasts. An increasing secretion of extracellular matrix can therefore stimulate the proliferation of endothelial cells, which is essential for angiogenesis and blood circulation in the skin microenvironment^23^. Since endothelial cells are the main source of vascular growth factors, they promote the formation of new blood vessels and improve blood supply and oxygen flow to the skin^24^. With all of these benefits that can cause a reduction in the appearance of skin wrinkles, retinol has become a star in the field of skin anti-aging.

Skin is the largest organ in the human body and the habitat of millions of commensal bacteria, known as the skin microbiome^25^. Skin microbiome plays an important part in protecting against pathogens, modulating immune response, and strengthening the skin barrier^26^. It is well-established that topical use of skin care products can alter the environment in which the skin microbiome lives, thereby altering the composition and function of the skin microbiome. Retinol, known for its antibacterial properties, has been shown to enhance host immune function in the skin^27,28^. A study by Harris et al^29^. demonstrated that dietary vitamin A can stimulate epidermal cells to produce antimicrobial protein RELMα, suggesting that endogenous vitamin A protects the skin against infection by reducing the colonization of pathogenic bacterial species. Additionally, there is growing evidence linking vitamin A deficiency with *Staphylococcus aureus* skin infections^30–32^. However, the effects of topically applied retinol on skin microorganisms have been poorly studied.

To fill the gap in this area, our study aims to investigate the impact of using retinol on the skin microbiome and understand the role of the skin microbiome in mediating the anti-aging properties of retinol. Using a multi-omics approach encompassing phenotypic, metagenomic, and metabolomic analyses, we seek to unravel the interactions between the host and its commensal microorganisms. Furthermore, we aim to identify microbial and metabolic biomarkers associated with anti-aging effects. The findings of our study can provide valuable insights from a microbiological perspective, guiding the development of next-generation anti-aging products. Ultimately, this research will contribute to a better understanding and utilization of the skin microbiome’s potential in anti-aging innovations.

## Materials and methods

### Study design and subject recruitment

Nine healthy Chinese women aged 22-38 years old were recruited, with an average age of 32.4, who presented with dulled and lifeless skin at baseline (Table S1). The participants agreed to refrain from using any cosmetics, medications, or supplements that could potentially affect the results. They demonstrated good cooperation throughout the study, maintained their regular lifestyle, and exhibited comprehension of the informed consent form as evidenced by their voluntary signature. In order to select suitable subjects, their skin condition was checked and a safety assessment was done according to Technical Code of Cosmetic Safety (2015 edition, China) by a dermatologist in advance.

Subjects were excluded from the study if they met any of the following conditions: 1) Presence of facial skin diseases that could influence the evaluation of test results; 2) High level of allergies; 3) Pregnancy, breastfeeding, or intent to become pregnant during the trial; 4) Severe cardiac, hepatic, or renal impairment, and severe immunocompromise; 5) Psychiatric disorders, severe endocrine disorders, or use of oral contraceptives; 6) Participation in drug clinical trials or other experiments within the past 30 days, or recent systematic use of drugs that could affect the test results; 7) Use of cosmetic products orally or topically within the past 2 weeks that could affect the test results; 8) Inability to cooperate with the test procedures; 9) Deemed unsuitable for participation in the study by the investigator’s judgment; 10) Experienced adverse reactions during the trial and deemed unsuitable for further participation by a dermatologist; 11) Other appropriate exclusion criteria.

A skin lotion containing 0.15% retinol developed by HBN (China) was used in this study. Subjects received the retinol lotion on Day 0, and it was checked and weighed at each follow-up visit. The retinol lotion was applied once a day at night for consecutive 28 days. Skin swabs and facial phenotype data were collected at five time points: day 0 (baseline), day 7, day 14, day 21, and day 28.

### Sample collection

Samples were collected at weekly intervals, with all procedures conducted in a controlled environment with regulated temperature and humidity (21±1℃, 50±10%RH). On the day of sample collection, each subject maintained their facial microenvironmental characteristics and refrained from washing their face for at least 12 hours.

Two adjacent sampling areas of approximately 2cm x 2cm were selected on the cheeks, designated as the “DNA sampling area” and the “metabolite sampling area”. Sterile cotton-tipped swabs (Winner, China) and saline (0.9% NaCl) were used to collect microbial DNA samples and metabolite samples, respectively, from the designated areas. The collected swabs were stored at −30°C. Once the facial swab collection was completed, subjects cleansed their face, dried their skin, and then rested for 30 minutes in the temperature- and humidity-controlled laboratory environment. Phenotype information of the participants was then gathered. Measurements were taken using the following instruments: Corneometer CM 825 (Courage + Khazaka Electronic GmbH, Cologne, Germany) for water content in the stratum corneum (WCSC), Vapometer for trans-epidermal water loss (TEWL), Skin-pH-Meter pH905 for skin surface pH, Sebumeter SM815 (Courage + Khazaka Electronic GmbH, Cologne, Germany) for sebum content. Additionally, facial image analysis was conducted using VISIA-CR (Canfield Scientific, USA) to measure porphyrin area, pore area, percentage of skin red zone area, and skin color values (L*, a*, and b*). Wrinkle indicators at the corners of the eyes, including wrinkle number, length, area, and volume, were assessed using the skin-fast optical imaging system PRIMOS-CR (Canfield Scientific, USA).

The recruitment, management of participants, sampling of the skin microbiome and metabolome, and skin measurements were conducted by Weipu Testing Technology Group Co., LTD (China) through a contract research arrangement. All participants provided written informed consent after fully comprehending the project’s details.

### Microbiome DNA extraction and library preparation

Genomic DNA samples from skin microbiome cotton swabs were extracted using DNeasy PowerSoil Pro Kit (Qiagen) according to the manufacturer’s instructions. Briefly, the swab samples were lysed via both lysis buffer and 15mins zirconium beads vortex. Crude lysate was then subjected to inhibitor removal for cleanup. Then, purified lysate was mixed with DNA binding solution and passed through a silica spin filter membrane. After a two-step washing regime, silica-bound DNA was eluted by 100μl 10Mm Tris elution buffer, and Qubit (Thermo Fisher Scientific) was used for DNA sample concentration determination. Extracted DNA samples were stored at −30°C until removed for library construction.

The metagenomics DNA libraries were constructed using VAHTS Universal Plus DNA Library Prep Kit for MGI (Vazyme) according to the manufacturer’s instructions. The single indexed paired-end libraries of genomic DNA were generated with an average size of 150bp using DNA from each sample. DNA fragmentation, adapter ligation and library amplification were carried out according to the kit protocol, and the amplification step was selected uniformly for 17 rounds of PCR. DNA libraries were purified using VAHTS DNA Clean Beads (Vazyme). The resulting libraries were sequenced by Geneplus on DNBSEQ-T7 (MGI Tech, China).

### Metagenomics sequencing data analysis

ATLAS^33^ v.2.16.3 pipeline was used to process metagenomic quality control, assembly, genomic binning and annotation. In brief, BBTools were used to remove PCR duplicates, reads were quality trimmed and contaminations from the PhiX and human genome were filtered out. Before assembly with MEGAHIT^34^, clean reads were error-corrected and merged. Assembled contigs were quality filtered and coverage rates were calculated using BBMap. Contigs were binned using MetaBAT2^35^ and MaxBin2^36^ and their predicted bins were combined by DAS-Tool^37^ to get metagenome assembled genomes (MAGs), which had at least 50% completeness and < 10% contamination based on the estimation by CheckM2^38^ and dereplicated by dRep^39^. Finally, 40 MAGs were clustered (95% average nucleotide identity). The open reading frame of each genome was predicted by Prodigal^40^, clustered by linclust^41^ to a nonredundant gene catalog and annotated by EggNOG and dram robustly. The phylogenetic tree of each MAG was constructed using GTDB and CheckM2, visualized in iTOL^42^ v6. The relative abundance of each representative MAG was calculated by dividing the number of reads that mapped to that MAG, corrected to the genome size and completeness, by the total number of reads in each sample.

The average raw reads number was 51.8±20.6M, after quality control and filtering host reads, the average microbial reads number was 16.5±13.2M, accounting for 34.2±24.8% of high-quality reads. Meanwhile, based on metagenomic clean reads, we used mOTUs3^43^(v.3.0.3 on Python v.3.8.12) to classify all reads into operational taxonomic units (OTU) clusters according to 97% similarity, taxonomy annotations were made based on the reference databases. HUMAnN3^44^(v.3.6) was used to quantify gene families based on metagenomic clean reads, definitions of gene families were provided by UniRef and KEGG databases. Gene pathway enrichment analysis was based on ReporterScore^45^, where reporter scores greater than 1.64 indicate significantly enriched pathways, the plus or minus sign represents the up- and down-regulated of the pathway compared to the control group (Day0, baseline).

### Genome-Scale Metabolic Model Construction

In this study, we constructed genome-scale metabolic models (GEMs) using metagenomic data derived from 9 samples. These samples were processed through Metagenome-Atlas analysis (v.2.16.3). This rigorous analytical process successfully yielded 40 Metagenome-Assembled Genomes (MAGs). The corresponding .fna files, encapsulating the genome sequences of these MAGs, served as the cornerstone for our GEM construction. To facilitate this construction, we employed gapseq^46^ (version 1.2), a robust and efficient tool renowned for its capability in constructing metabolic pathways and identifying transport proteins via homology-based methods. A pivotal component of our methodology was the completion of the ‘gapseq doall’ process. This comprehensive step involved the identification of metabolic pathways (using the ‘find’ function) and transport proteins (using the ‘find-transport’ function), all discerned through homology. This was followed by the creation of preliminary draft models and further refinement of these models through a meticulous gap-filling process (using the ‘fill’ function), effectively addressing and rectifying any metabolic gaps present.

### Metabolic Network Analysis

For the analysis and manipulation of the reconstructed metabolic models, we utilized the COBRA Toolbox^47^ (2024 release). This MATLAB toolbox is integral for constraint-based modeling of metabolic networks. Linear and integer programming simulations within these networks were executed using the Gurobi Optimizer (version 1003). The MetaboReport tool within the COBRA Toolbox was employed for quality control and reporting.

Flux Balance Analysis (FBA) for each reaction in the models was performed using the optimizeCbModel function in the COBRA Toolbox. This function optimizes metabolic networks based on specified objectives, providing insights into the metabolic capabilities of our samples. To gain a clearer understanding of the influence of metabolites and microbes on the reactions of interest, we computed the shadow prices using the FBAsolution.f and FBAsolution.y functions. These functions are part of the standard COBRA Toolbox functionality and are used for calculating objective values and shadow prices, respectively.

### LC-MS metabolomic data collection and analyses

The experiment and preprocessing of non-targeted metabolomic data were conducted by Shanghai OE Biotech Co., Ltd (China). Briefly, Two skin swab samples were taken in a 5 mL centrifuge tube and mixed with 1 mL of pre-cooled methanol-water (V:V=4:1, Thermo Fisher), after brief sonication in ice water for 20 min, all samples were left at −40℃ overnight. The samples were then centrifuged for 10 min(12,000 rpm, 4°C), and 500 μL of which was vented into an LC-MS injection vial. Then, the samples were re-dissolved in 300 μL of methanol-water (V:V=1:4, containing mixed internal standard, 4 μg/mL), vortexed for 30 s, sonicated for 3 min in an ice-water bath, and placed in −40 °C for 2 h. Then, the samples were centrifuged for 10 min(12,000 rpm, 4° C), 150 μL of the sample supernatant was aspirated with a syringe, filtered using a 0.22 μm organic-phase pinhole filter, transferred to an LC injection vial, and stored at −80 °C until LC-MS analysis. For quality control (QC) sample preparation, equal volumes of extracts from all samples were mixed. For LC-MS metabolomic data collection, all extracted skin swab samples were analyzed by a Waters ACQUITY UPLC I-Class plus/Thermo QE plus instrument, coupled with ACQUITY UPLC HSS T3 (100 mm×2.1 mm, 1.8 μm) column.

Prior to pattern recognition, data was preprocessed. Raw data were subjected to baseline filtering, peak identification, integration, retention time correction, peak alignment and normalization by the metabolomics processing software, Progenesis QI v3.0 software (Nonlinear Dynamics, Newcastle, UK). Metabolites were identified based on multiple dimensions such as RT (retention time), exact mass number, secondary fragmentation, and isotopic distribution, and were analyzed using The Human Metabolome Database (HMDB), Lipidmaps (v2.3), and METLIN databases, as well as the LuMet-Animal local database. The data matrix was further processed by removing the peaks with missing values in more than 50% of the samples and substituting the remaining missing values with half of the minimum value. The compounds obtained from the characterization were screened based on a score of 36 out of 80 for the results of the characterization of the compounds, with a score of 36 or less being regarded as an inaccurate characterization result and deleted. Score descriptions are as follows: 80 in total, the exact molecular weight match of the primary mass spectrum (20 points), the fragmentation match of the secondary mass spectrum (20 points), the isotopic distribution match (20 points), and the retention time match (20 points), and in general, it is considered that the higher the score, the more accurate the characterization will be.

Additionally, metabolic pathway analysis was performed using the web-based MetOrigin^48^ (https://metorigin.met-bioinformatics.cn/home/), with its databases identifying where metabolites come from: host, bacteria, or both.

### Statistical Analysis and Data Visualization

Statistical analysis and data visualization were performed by using Microsoft Excel (Microsoft Inc., Redmond, WA, United States) and R software version RStudio v4.3.2(R Foundation for Statistical Computing, Vienna, Austria) with the ggplot2 library (version 3.4.4). The differential abundance of bacterial taxa, gene and metabolites and alpha diversity indexes between different time points and baseline was calculated by the Wilcoxon rank-sum test. The beta diversity was determined by Bray-Curtis distance, and the difference between different time points and baseline was calculated by the Adonis Test. Spearman’s and Pearson’s correlation analysis was carried out to determine the relationship between the skin microbiota, skin metabolites, microbial genes and skin phenotype. The flux consistency and stoichiometric consistency of the models were evaluated and compared, providing an effective means for visual representation of the analytical results. The compilation of all shadow price analysis results was carried out using Python (version 3.8.8) with the Pandas library (version 1.2.4). For the visualization of metabolic networks, we utilized the createMetIntrcNetwork function from the COBRA Toolbox. This function allows for the creation of interactive metabolic network graphs, highlighting the interconnections and flux distributions within the network. The parameters such as ‘fluxes’, ‘threshold’, and ‘excNodesWithDeg’ within this function were adjusted to suit the specifics of our analysis, offering a detailed and insightful representation of the metabolic interactions. The p-value was corrected by the Benjamini & Hochberg (BH) method, and p < 0.05, FDR<1 was considered significantly altered unless otherwise stated.

## Results

### 1. Retinol alleviates aging related conditions of human skin

The skin phenotypes of each individual enrolled were collected at five time points. Principal coordinates analysis (PCoA) revealed that participants experienced an overall change in skin phenotype after application of retinol. The Adonis test demonstrated significant separation from Day 0 for Day 21 (adj.p=0.010) and Day 28 (adj.p=0.017), indicating an altered phenotype compared to the baseline (Fig.S1A). Moreover, more than half (eight out of fourteen) of the phenotypic measurements, including water content in the stratum corneum (WCSC), trans-epidermal water loss (TEWL), pH value, percentage of the red area, and various wrinkle parameters (number, length, area, and volume), exhibited significant changes compared to the baseline (Fig.1).

**Figure 1.**
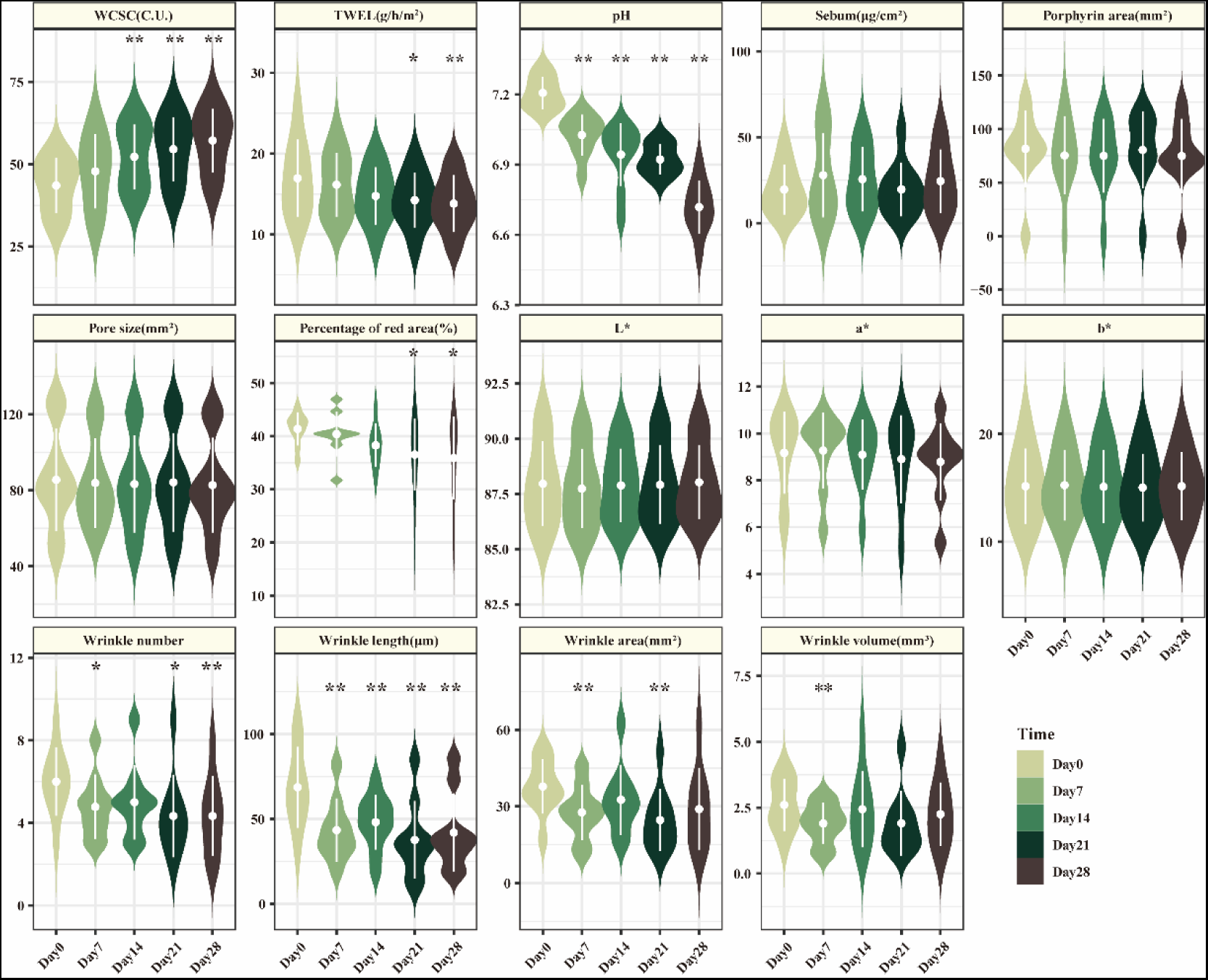
The temporal variation of skin phenotypic traits. Statistical significance levels of data relative to baseline changes were assessed using the Wilcoxon paired test and adjusted using the Benjamini & Hochberg (BH) method. Displayed results are adjusted p values. Unmarked indicates non-significance, * indicates adj.p ≤ 0.1, ** indicates adj.p ≤ 0.05. Data are shown in Table S2.

The topical use of the retinol resulted in enhanced water retention capability of the skin, with a 31.4% increase in WCSC (adj.p=0.050, Wilcoxon paired test) and an 18.4% decrease in TEWL at day 28 (adj.p=0.004, adj.p=0.050, Wilcoxon paired test), respectively. Notably, WCSC on day 28 showed a significant increase not only compared to the baseline but also compared to days 7 and 14 (adj.p=0.004, adj.p=0.050, Wilcoxon paired test), suggesting that the stratum corneum’s water retention ability improves with prolonged retinol use. The decrease in TWEL indicated that retinol markedly improved and repaired the skin barrier, helping to retain skin moisture and prevent water loss. Retinol also demonstrated its effectiveness in sedative and anti-inflammatory skincare properties, as evidenced by a significant reduction in the size of red areas: a decrease of 11.6% on day 21 and 13.2% on day 28 compared to the baseline level (Table S2).

The pH values of the skin cheek surface exhibited a decline from 7.21 (average value on Day 0) to 6.72 (average value on Day 28) while using retinol, indicating the gradual formation of a weakly acidic environment on the facial skin, suggesting that retinol can modulate and maintain acid-base balance of skin.

Meanwhile, multiple wrinkle-related indicators, including the wrinkle number, length, area, and volume at the corners of the eyes, displayed significant reductions starting from Day 7 compared to the pre-retinol use conditions (Table S2). Specifically, the wrinkle number decreased significantly on Day 7, 21 and 28, with the most substantial reduction observed on Day 21 and 28, resulting in a 27.8% decrease relative to baseline. The wrinkle length significantly decreased at all four sampling time points compared to baseline. By Day 28, the average wrinkle length decreased from 68.7μm to 42μm, representing a reduction of 38.8% compared to baseline. Wrinkle area exhibited significant reductions on Day 7 and 21, with a notable decrease of 34.8% observed on Day 21. Finally, the wrinkle volume exhibited a significant decrease of 26.9% at Day 7. These findings highlight the potent anti-aging properties of retinol and its efficacy in improving facial wrinkle conditions.

### 2. Retinol reshapes human skin microbiome microecology

The application of the retinol had a dramatic impact on the restructuring of skin microbiome microecology (Fig.S1C). Species-level alpha diversity (Shannon diversity index and species evenness index) was significantly lower on day 7 compared to day 0 (Fig.2B, p=0.031, paired Wilcoxon test). This decrease in diversity could be attributed to an imbalance in the relative distribution of certain species within the microbial community. Specifically, there was a significant decrease in the relative abundance of *Corynebacterium accolens*, a skin bacterium ranked among the top 20 abundant species, on day 7 compared to the baseline (Fig. S1B). This reduction may have allowed other species to occupy a relatively larger ecological niche, resulting in a decline in microbial diversity. Notably, opportunistic pathogens such as *Stenotrophomonas maltophilia*, *Acinetobacter johnsonii*, *Pseudomonas sp.*, and *Sphingomonas hankookensis* showed significant decreases in their relative abundances at three consecutive time points compared to the baseline (Fig.2C). This suggests that the retinol-containing skincare product possesses antimicrobial properties and can reduce the colonization of pathogenic bacteria on the skin surface. We also noticed an increase in the relative abundance of *Neisseriales species incertae sedis* (Fig.S1B) and *Corynebacterium jeddahense* (Fig.2C). However, their specific functions remain unclear.

**Figure 2.**
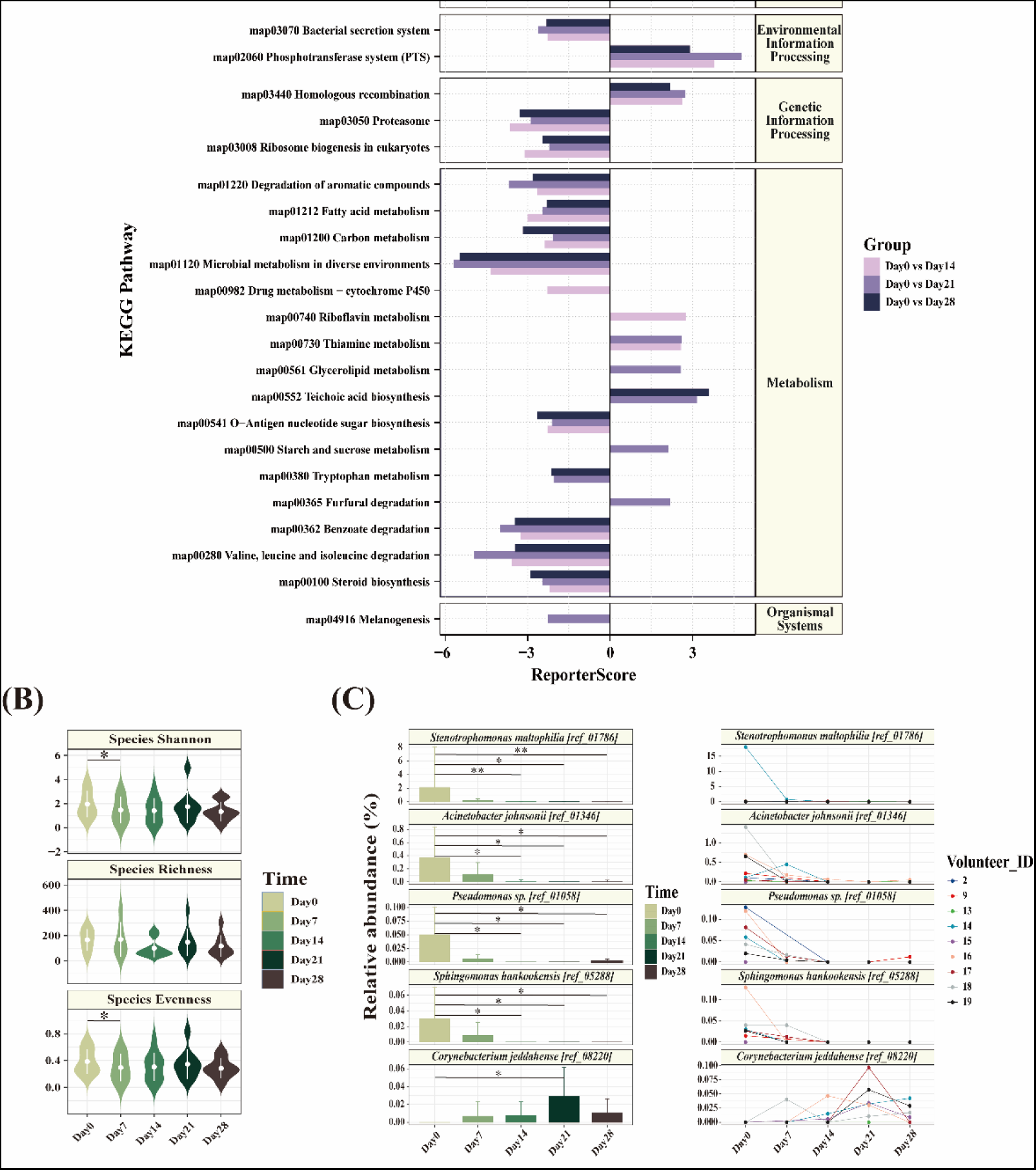
Effects of retinol on the structure of skin microbiome. (A) Gene pathway enrichment analysis based on ReporterScore. A ReporterScore with an absolute value greater than 1.64 indicates significant enrichment of the gene pathway, with positive or negative signs denoting upregulation or downregulation compared to the control group (Day 0, baseline). (B) Skin microbiome Alpha diversity indexes. (C) Inter-group differential species abundance changes. Statistical significance levels of data relative to baseline changes were assessed using the Wilcoxon paired test. Displayed results are p values. Unmarked indicates non-significance, * indicates p ≤ 0.05, ** indicates p≤ 0.01.

Retinol also exerts an impact on the functionality of the skin microbiome. Starting from Day 14, several microbial gene pathways displayed altered regulation levels (Fig. 2A). Notably, the thiamine (Vitamin B1) metabolism gene pathway was enriched on both Day 14 and Day 21, marked by a peak in the abundance of a thiamine metabolite, biotin thiamine, on Day 14, which significantly increased compared to Day 0 (Fig. S2A). Metabolomic data also showed that the microbial and host thiamine metabolic pathway was significantly (p=0.026) enriched on Day 14, marked by increased intensities of pyruvic acid and L-Tyrosine (Fig. S2A). All the three substances are biosynthetic precursors of thiamin. Thiamin has a variety of benefits for the skin, including increasing the expression of collagen, promoting skin cell growth and repair, and maintaining skin elasticity. Our findings suggest that retinol helps to promote the synthesis and utilization of thiamine by the skin microbiome, thereby enhancing the function and health of the skin barrier. Furthermore, the riboflavin metabolism pathway showed enrichment on Day 14 (Fig. 2A), accompanied by a notable decrease in the abundance of riboflavin on Day 14, while the intensity of riboflavin 5’-phosphate sodium, a bio-active form of riboflavin, exhibited an increase relative to baseline on Day 21 and Day 28(Fig. S2A). Riboflavin phosphate sodium salt form is an essential micronutrient, and plays an important role in the health of the skin, mucous membranes, and eyes. Retinol makes microorganisms more inclined to utilize or convert riboflavin into the active form, riboflavin 5’-phosphate sodium. Additionally, certain gene pathways demonstrated decreased expression levels, including biofilm formation of *Vibrio cholerae* and flagellar assembly were down-regulated on Day 21. And the bacterial secretion system and O-antigen nucleotide sugar biosynthesis, both were consistently down-regulated during the last three time points (Fig. 2A). The down-regulation implies that retinol may possess antimicrobial and anti-inflammatory capabilities by inhibiting bacterial metabolic activity, secretion of bacterial products, and influencing the integrity of bacterial structure.

### 3. Retinol stimulates skin microbiota’s secretion of diverse beneficial metabolites for synergistic anti-aging effects

We further utilized MetOrigin to perform tracing analysis of metabolites (based on the databases of MetOrigin tracking to determine whether they originated from the host, microorganisms, or co-metabolism) and metabolic pathway enrichment analysis. Of note, nicotinate and nicotinamide metabolism pathway (hsa00760) enriched in the host on Day 21 (Fig. S3A), and supported by increment of N1-methyl-4-pyridone-3-carboxamide which is associated with this pathway (Fig. S2B).

In the microbe, degradation of flavonoids (ko00946), phenylalanine, tyrosine and tryptophan biosynthesis (ko00400), and biosynthesis of various plant secondary metabolites (ko00999) were up-regulated (Fig. 3A). Based on the databases of MetOrigin, we found a variety of microbial-origin metabolites significantly related to the enrichment of the above pathways, some of which have been reported or experimentally verified to be beneficial to the skin. Maesopsin and apigenin were related to ko00946, where apigenin was known as an anti-tumor substance that is particularly helpful in preventing and reversing the formation of abnormal skin^49–52^. Quinic acid, 3-dehydroquinic acid and protocatechuic acid were related to ko00400, where quinic acid was reported to have an antiphotoaging effect by protecting human dermal fibroblasts^53,54^ and protocatechuic acid was demonstrated to have anti-oxidate and anti-aging effects by inducing dermal fibroblasts to synthesis type-1 collagen^55–57^. In ko00999, (+)-pinoresinol and secoisolariciresinol were significantly related, the former was reported to stimulate keratinocyte proliferation^58,59^ and the latter was reported to suppress atopic dermatitis in the mouse when administered orally^60^. (Fig. 3B-D)

**Figure 3.**
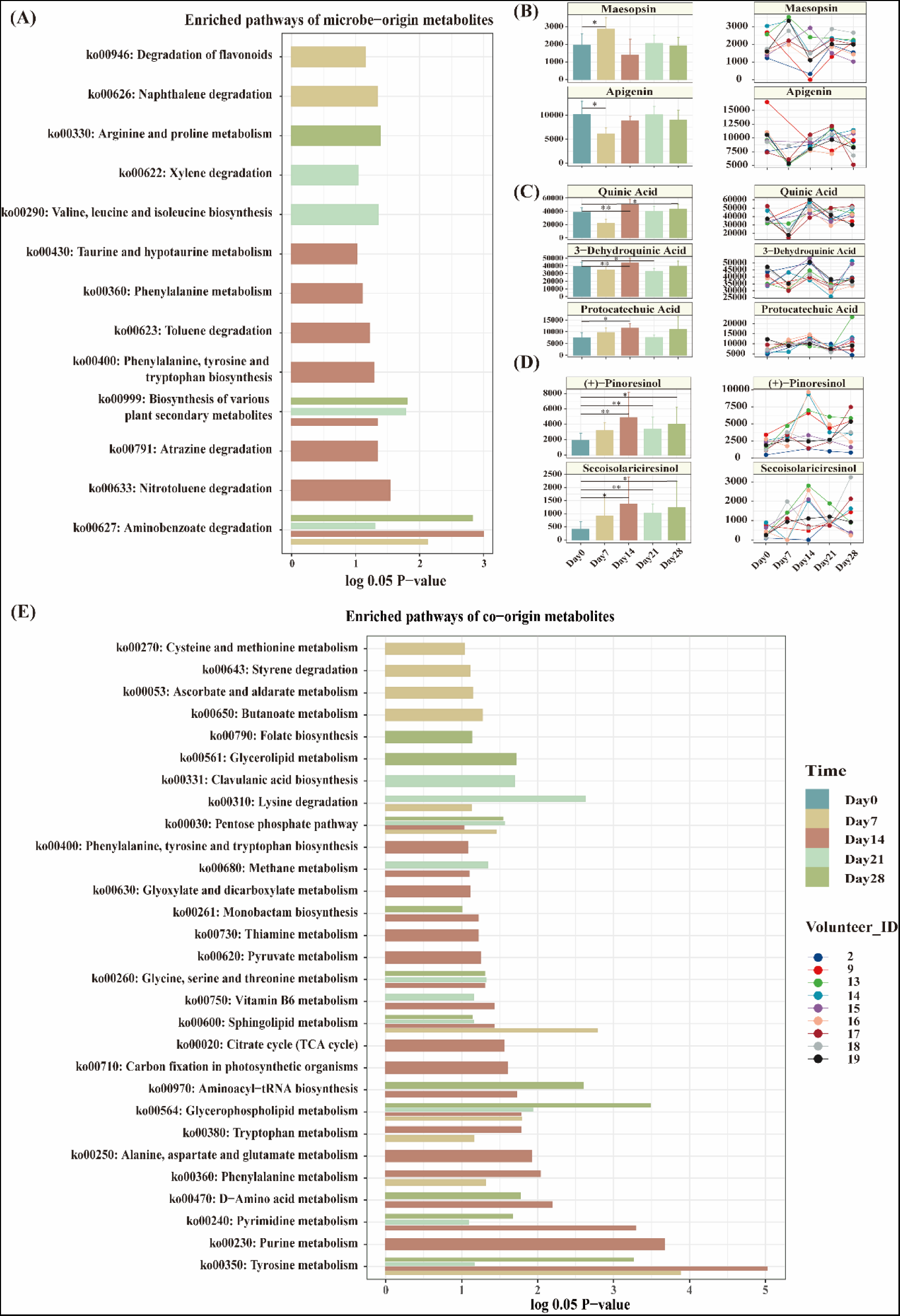
Metabolic pathway enrichment analysis based on MetOrigin. (A) and (E) enriched pathways of microbe and co-origin metabolites compared to baseline. Relative abundance changes of metabolites significantly associated with (B) degradation of flavonoids, (C) phenylalanine, tyrosine and tryptophan biosynthesis and (D) biosynthesis of various plant secondary metabolites.

For co-origin metabolic pathways, we focused specifically on the following three enriched pathways: thiamine metabolism (ko00730), vitamin B6 metabolism (ko00750), and tyrosine metabolism(ko00350) (Fig. 3E). Thiamine metabolism was enriched on Day14, which is consistent with the timing of microbial thiamine metabolism gene pathway enrichment (Fig. 2A). This indicates that skin microbes play an important role in thiamine metabolism and the function was enhanced by the application of retinol. Three metabolites were found to be related to vitamin B6 metabolism, including L-glutamine, pyridoxine and pyridoxamine (Fig. S2C). L-glutamine is an important skincare ingredient, it helps the body to produce anti-oxidants, thus providing ROS protection^61,62^. Pyridoxine has the potential to prevent pigmentation and stimulate filaggrin production in human epidermal keratinocytes^63,64^. At last, tyrosine metabolism is associated with melanogenesis, with dopaquinone and 5,6-dihydroxyindole (Fig. S2D) related to this pathway. The abundance of the two metabolites increased on Day 7 and Day 28, respectively.

### 4. The formation of an acidic skin is associated with the secretion of acidic metabolites by skin microorganisms

The use of retinol has been demonstrated to induce significant changes in the skin phenotype, including a reduction in skin pH levels, from 7.21 to 6.72. Past research has demonstrated that skin pH largely influences skin barrier homeostasis, stratum corneum integrity and cohesion, and antimicrobial defense mechanisms^65^. Lower pH contributes to the activity of key enzymes of permeability barrier synthesis, such as β-glucocerebrosidase and acidic sphingomyelinase^66^. Most retinoids have optimum stability at a pH of 6-7. The pH value of the retinol lotion used in this study was 6.42, which is lower than the average baseline skin pH of the subjects. However, in addition to the acidity of the lotion itself, we have found that skin microbes also play a key role in the pH drop process. The Mantel test showed a significant positive correlation between dermal microbial species and the skin pH of the subjects (Fig. 4A). Six species with a decreasing relative abundance, namely *Cutibacterium modestume, Sphingomonas hankookensis, Acinebacter johnsonii, Pseudomonas sp.* (Fig. 4B), *Stenotrophomonas maltophilia* and *Braychybacterium paraconglomeratum* (Table S6, S7) were found to be positively correlated with pH. Whereas one microorganism, *Cornebacterium jeddahense*, with an increased relative abundance, is significantly negatively correlated with skin pH (adj.p=0.047). On one hand, the change reflects the ecological adaptations and differential metabolic characteristics of skin microorganisms. On the other hand, the modulation of host skin pH by microorganisms could be associated with the secretion of microbial metabolites. A variety of metabolites classified by MetOrigin as microbe-origin were acid, including (R)-mandelic acid, phenylglyoxylic acid, protocatechuic acid, vanillic acid and syringic acid, among others (Table S5). Of the above metabolites phenylglyoxylic acid was significantly negatively correlated with skin pH (adj.p=0.023, Table S6, S7). Phenylglyoxylic acid was positively related to *Neisseriales species incertae sedis* [meta_mOTU_v3_13131], and may be a source of nutrients or metabolites for the microorganism (Table S8, S9).

**Figure 4.**
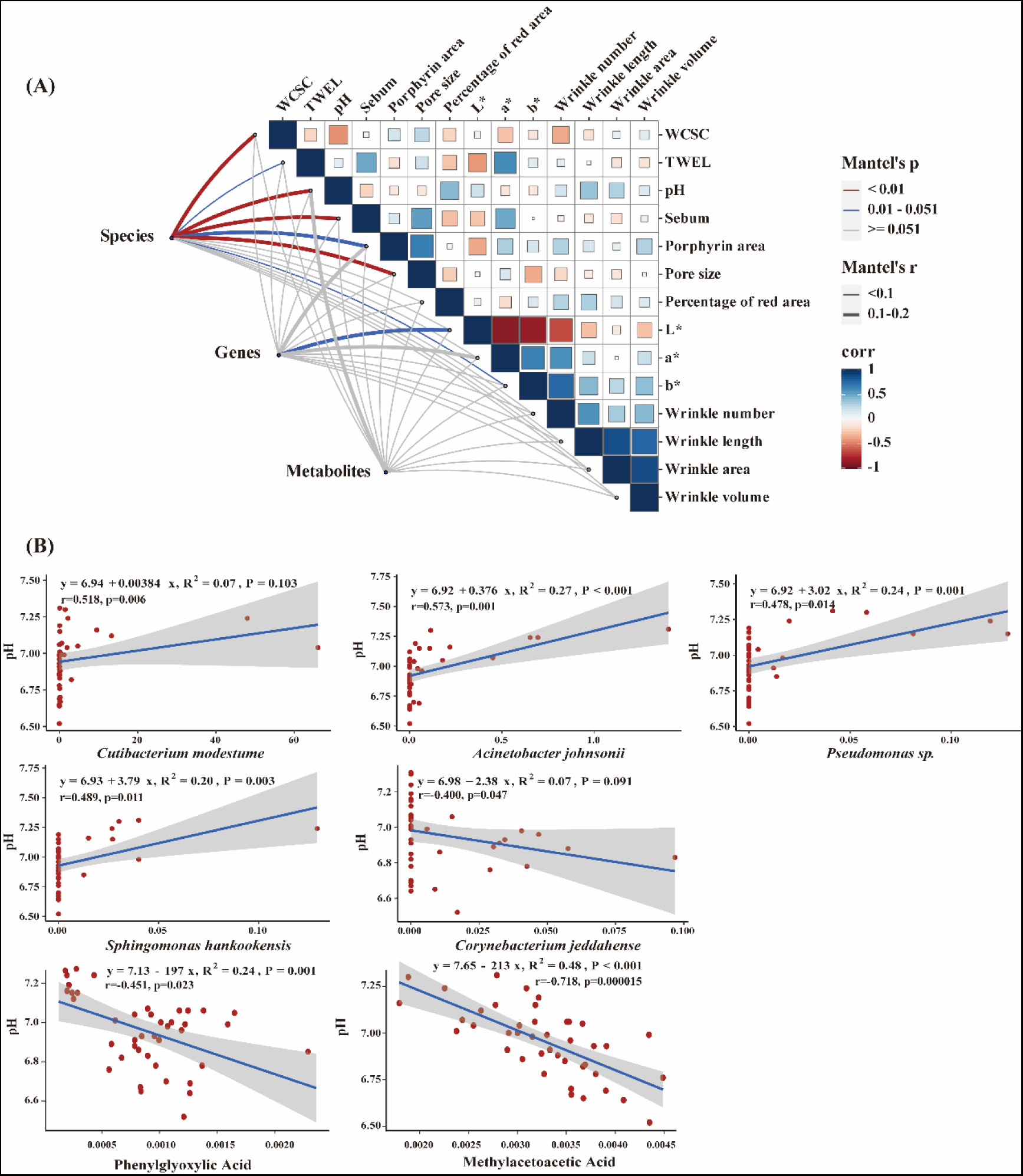
Multi-omics data integration and its correlation analysis with skin phenotypes. (A) Mantel matrix correlation analysis, the thickness of the lines represents the correlation coefficient (r value) between matrices, with thin lines indicating correlation coefficients <0.1 (negative correlation), thick lines indicating correlation coefficients between 0.1 and 0.2 (positive correlation). The color of the lines represents the significance level (p-value) of the correlation between matrices, where red indicates p <0.01, representing a strong significant correlation, blue indicates 0.01 <p <0.05, representing a significant correlation, and gray indicates p≥0.051, representing no significant correlation. “Corr” indicates the Pearson’s self-significant correlation of each phenotype indicator, with blue representing positive correlation and red representing negative correlation. (B) Spearman’s correlation analysis between skin microbial species and skin metabolites with skin pH. Data are shown in Table S6 and S7.

### 5. The combined anti-aging effects of skin microorganisms and retinol

The intensity of a soluble metabolite of retinol, 1-O-all-trans-retinoyl-β-glucuronic-acid (RAG), was found to reach its peak on Day 7 and remain at baseline level on Day 14, 21 and 28 (Fig. 5A). The presence of RAG can be a mixed retinol metabolism result by both human body and skin microorganisms. Pearson’s correlation analysis revealed significant positive associations between RAG and 11 microorganisms, including 3 *Corynebacterium* species and *Lactococcus lactis* (Fig. 5B, Table S10, S11).

**Figure 5.**
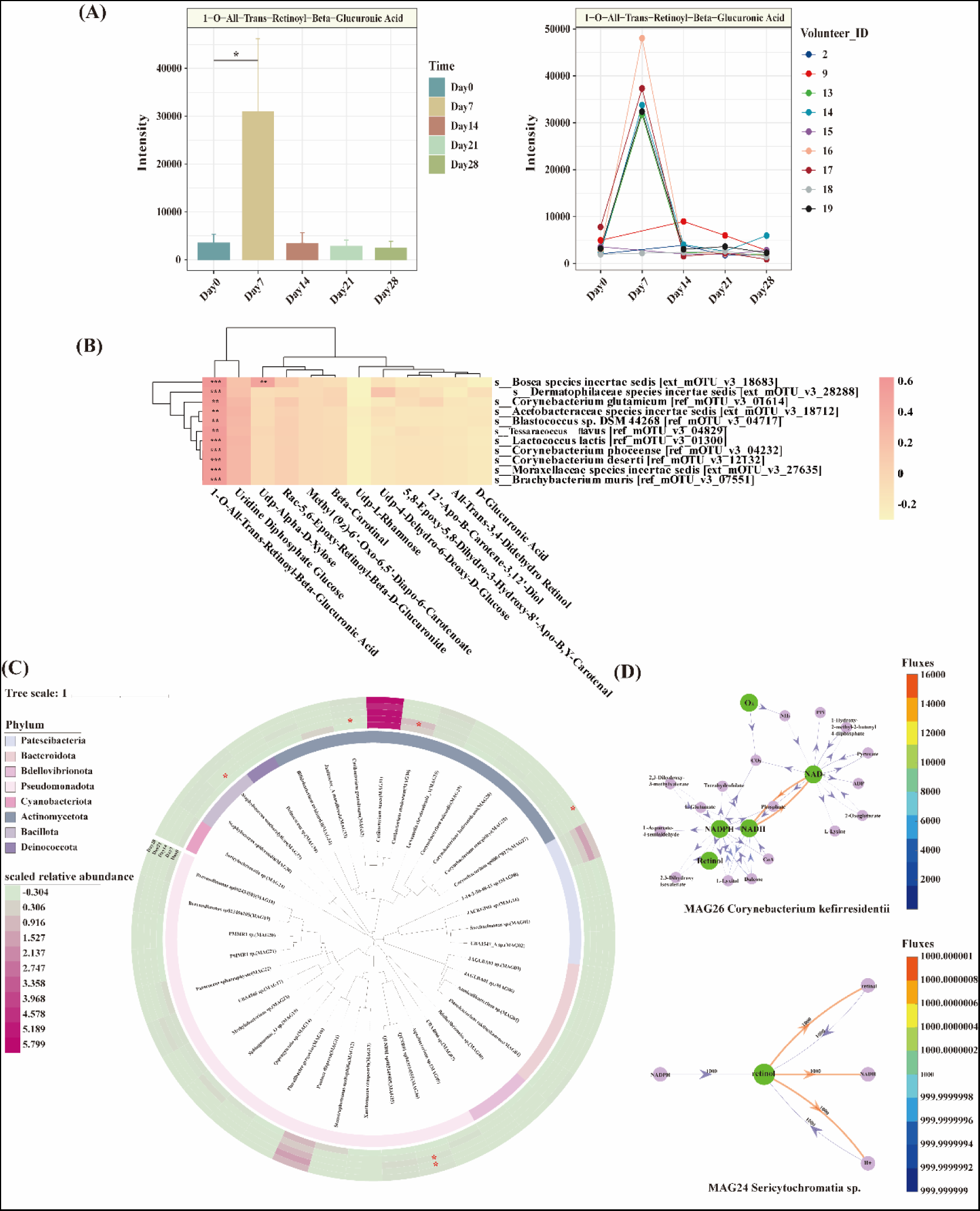
Skin microbiota participate in retinol metabolism through both direct and indirect ways. (A) The intensity changes of a soluble retinol metabolite, RAG. (B) Correlation heatmap derived from Pearson’s correlation analysis showing representative microbial associations related to retinol metabolism, full data in ST10 and 11. The color depth indicates the magnitude of correlation coefficients. Displayed results are adjusted p values (adj.p values). Unmarked indicates non-significance, * indicates 0.05 < adj.p ≤ 0.1, ** indicates 0.01 < adj.p ≤ 0.05, *** indicates adj.p ≤ 0.01. (C) Phylogenetic tree of 40 MAGs species. Displayed results are adjusted p values (adj.p values). Unmarked indicates non-significance, * indicates 0.05 < adj.p ≤ 0.1. (D) The metabolic reaction flux networks of MAG24 and MAG26 involving in retinol metabolism.

To further explore the involvement of skin microorganisms in retinol metabolism, we constructed 40 metagenome-assembled genomes (MAGs) and reconstructed their metabolic networks at the genome scale (Fig. 5C). Reaction flux values were calculated for all reactions related to retinol metabolism (map00830). MAG24, identified as *Sericytochromatia sp.*, and MAG26, identified as *Corynebacterium kefirresidentii*, exhibited high reaction fluxes directly involved in retinol metabolism. Specifically, they were responsible for oxidizing retinol to retinaldehyde, which represents the initial step in the retinol metabolism pathway (Fig. 5D).

15 out of 40 MAGs were identified to be able to consume UDP-glucose(R00286) at high reaction fluxes, including *Pluralibacter gergoviae (MAG10)*, *Pantoea dispersa (MAG11)*, *Sericytochromatia sp. (MAG24) and Corynebacterium kefirresidentii (MAG26)*, etc (Table S12). UDP-glucose is a carbon source for microorganisms and also a hydrolysis product of RAG. We further hypothesized that skin microorganisms can be involved in retinol metabolism indirectly through the production and consumption of UDP-glucose. Since RAG can be hydrolyzed to retinoic acid and UDP-glucuronate (R02902), and the latter can be hydrolyzed to UDP-glucose (R00286), thus RAG hydrolysis is facilitated by the continuous use and consumption of UDP-glucose by microorganisms.

## Discussion

In this study, we focus on how skin microbiome mediates the anti-aging effects of retinol, and the interactions among retinol, skin microbes and metabolites in the skin microenvironment. As retinol is an effective anti-aging agent widely used in skin care products, however, whether its anti-aging efficacy is associated with skin microbes and how microbes mediate its anti-aging effects remains unexplored. By integrating multi-omics data from skin traits, metagenomics and metabolomics data, we found that the retinol has a strong anti-aging effect, with visible improvement in several skin epidermal wrinkle indicators.

Vitamin B comprises a group of water-soluble vitamins, including thiamine (VB1), riboflavin (VB2), niacin or nicotinamide (VB3), pyridoxine (VB6), etc. The synthesis and metabolism of vitamin B are essential for skin health. Our study reveals that retinol significantly influences the synthesis and metabolism in the skin by skin microbiota. Following retinol application, the gene pathway of thiamine (VB1) metabolism of skin microbiota was significantly enriched on Day 14 and 21, while riboflavin (VB2) metabolism pathway was significantly enriched on Day 21. The niacin and nicotinamide (VB3) metabolism metabolic pathway of the host was significantly enriched on Day 21, and the co-metabolism pathways of thiamine (VB1) and pyridoxine (VB6) between the host and microbiota were enriched on Day 14 and Day 14 and 21, respectively. These findings suggest that retinol application alters the metabolic pathways of vitamin B synthesis and utilization in both the host and skin microbiota, promoting the growth and proliferation of skin microbiota with more abundant vitamin B metabolism genes and increasing the secretion of vitamin B-related metabolites by skin microbiota.

Through MetOrigin analysis of metabolites significantly associated with enriched metabolic pathways such as vitamin B metabolism and flavonoid metabolism, we found that retinol use promoted the secretion of probiotic metabolites by skin microbes with synergistic anti-aging effects. These include apigenin, quinic acid, protocatechuic acid and (+)-pinoresinol, which possess antioxidant, anti-inflammatory and anti-aging properties. Our study also screened out several microbial metabolites with potential skin probiotic value, including maesopsin, 3-dehydroquinic acid and secoisolariciresinol. The skin anti-aging efficacy of these microbial metabolic substances awaits further validation through in vitro experiments.

In addition to this, retinol has the effect of lowering the acidic pH of the skin, which is contributed by two components, the acidity of the lotion itself and the acidic metabolites secreted by the skin microbiota that is restructured with retinol. Over the years, many *in vitro* and *in vivo* studies have focused on the effects of pH changes on skin microorganisms^65,67,68^, e.g., a physiologically acidic pH on the skin surface reduces the growth and adhesion of pathogenic bacteria and supports the production of antimicrobial peptides, but conversely, microbial shaping of the skin’s acidic environment is understudied. Traced back through MetOrigin, we identified acidic metabolites of significantly increased abundance from a variety of microbial sources, and Spearman’s correlation analyses also revealed microbial species associated with decreasing pH. We, therefore, proposed that the use of retinol altered the structure and composition of the skin microbiome, and that the remodeled microbiome secretes more acidic metabolites, such as phenylglyoxylic acid, which reduced skin pH. There has been a great deal of research^69,70^ showing that low pH contributes to the protective functions of the skin, including permeability barrier homeostasis^71,72^, stratum corneum stability^66,73,74^, and epidermal antimicrobial defenses^67,75,76^. Meanwhile, it has been reported^77–80^ that the skin of the aging population has a higher pH relative to the skin of the younger population, and that the skin barrier of the elderly is more easily disrupted and slower to repair than the skin barrier of the younger population. Lower pH reduces the enzymatic activity of cutaneous desquamation-related kallikrein-related peptidases and increases the activity of two ceramide and sphingomyelin lipid-processing enzymes, β-glucocerebrosidase and acid sphingomyelinase, whose optimal enzyme activity is acidic, thereby enhancing stratum corneum recovery and regeneration and protecting the skin barrier. Evidenced by an increase in WCSC and a decrease in TEWL, which reflect the elevated role of the lipid composition of ceramides, cholesterol, and free fatty acids within the stratum corneum in impeding the rate of water loss and the transdermal penetration of external harmful substances, we found that the skin barrier was indeed enhanced in subjects after the application of retinol. Thus, skin microbes play an important role in the anti-aging process of retinol by regulating skin pH acidity level.

Due to the inherent antimicrobial and antioxidant properties of retinol, the use of retinol led to structural and functional changes in the skin microbiome. The evolution of the skin microbiota has led to the emergence or enrichment of symbiotic bacteria adapted to retinol metabolism. The skin microbiota involvement in retinol metabolism has rarely been reported yet. Supported by Spearman’s correlation analysis result, we revealed that *Corynebacterium* genus play an important role in retinol metabolism. By genome-wide metabolic network analyses of the 40 MAGs constructed, we found two skin flora, *Sericytochromatia sp.* and *Corynebacterium kefirresidentii* can potentially contribute to the first step of retinol metabolism, oxidizing retionol to retinaldehyde. To function in the human body, retinol must be oxidized to retinoic acid, which is the bioactive form of retinol and retinaldehyde and binds to receptors to exert its effects^13,81^. The oxidation of retinol to retinaldehyde is an indispensable step in the synthesis of retinoic acid *in vivo*^10^. Thus, skin microorganisms increase the effectiveness and bioavailability of retinol in the skin layer and accelerate the production of retinoic acid.

RAG is an all-trans retinoic acid and glucuronic acid conjugate, first discovered in 1964. Its formation is catalyzed by UDP-glucuronosyltransferase, which increases the water solubility and inactivate retinoic acid by endogenous or exogenous glucuronidation. Therefore, RAG is generally considered to be an excretion product of retinoids in gastrointestinal^82^. It is reported that gut microorganisms can remove glucuronic acid as a carbon source by expressing β-glucuronidase enzymes, effectively reversing the inactivation in mammals^83^. Accordingly, we proposed another possible way by which microorganisms may be involved in retinol metabolism, by promoting RAG hydrolysis through the consumption of UDP-glucose, which in turn prolongs the duration of action of retinol on the skin surface.

The sample size in the current study is relatively small while future research should be conducted in larger populations to further discuss and validate our results. Overall, our findings demonstrate the anti-aging effects by topical application of retinol, and skin microbiota exert combined anti-aging effects with retinol by shaping the acidic environment and participating in retinol metabolism.

## Acknowledgement

This work is supported by Scientific Research Start-up Funds (QD2021005N), and Shenzhen Science and Technology Program (WDZC20220819134430002).

