## Supplementary Figures for "Decoding the anti-aging effect of retinol in reshaping the human skin microbiome niches"

Figure S1.

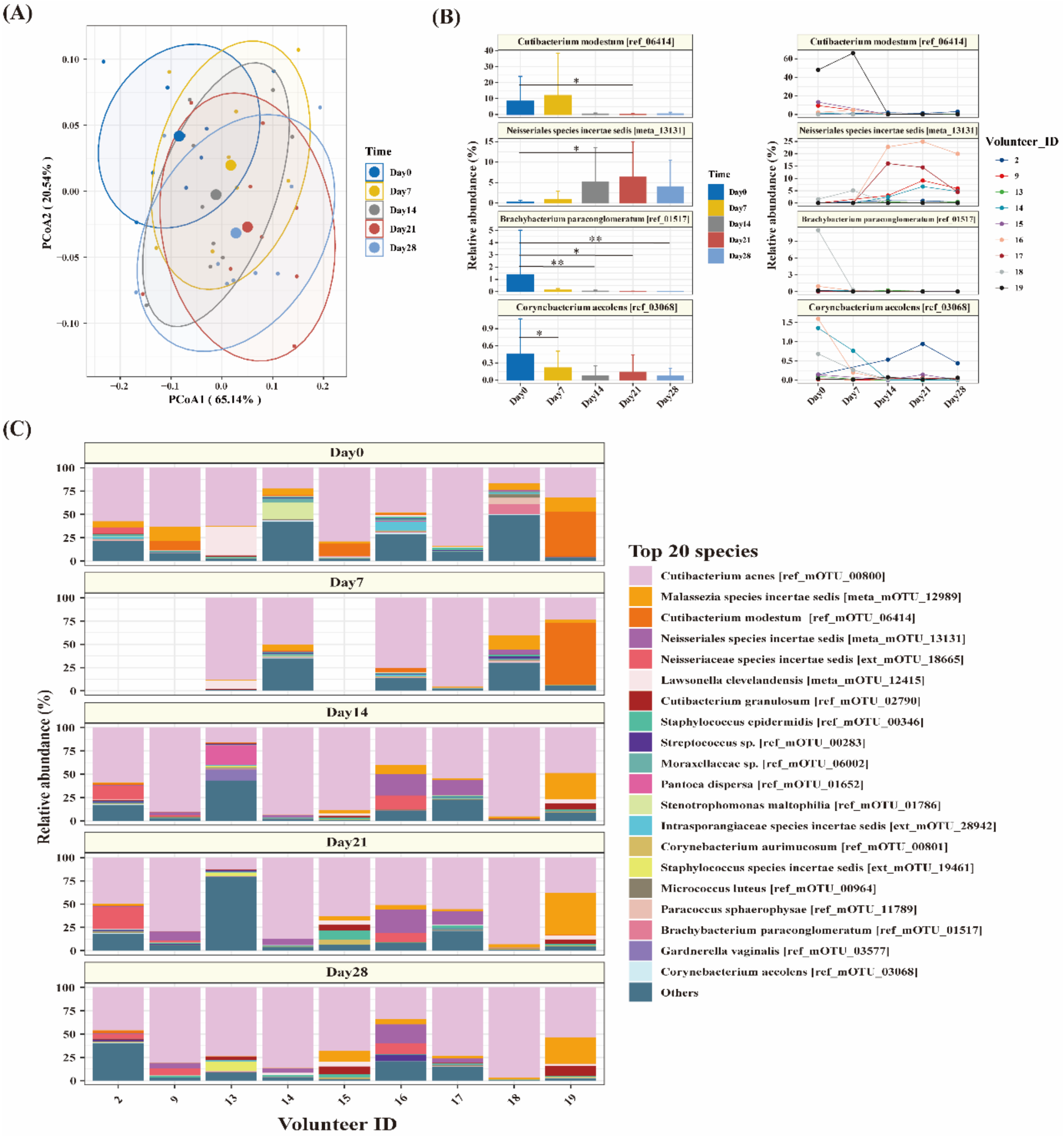


(A) The Principal Coordinates Analysis (PCoA) plot illustrates the overall differences in phenotypic indicators that show significant changes relative to the baseline following the use of retinol. PC1 and PC2 account for 65.1% and 20.5% of the total variance in the data, respectively. A distinct separation between Day 21 and Day 28 from Day 0 is observed, indicating significant phenotypic changes in the subjects. (Adonis Test). (B) The top 20 skin microorganisms that exhibit significant changes in relative abundance compared to the baseline. Statistical significance levels of data relative to baseline changes were assessed using the Wilcoxon paired test. Displayed results are p values. Unmarked indicates non-significance, * indicates p ≤ 0.05, ** indicates p≤ 0.01. (C) Distribution of the top 20 skin microbial species in each sample based on their relative abundance. The top 20 species were ranked according to their average abundance across all samples. Samples 2, 9, and 15 on day 7 were excluded due to failed DNA library preparation.

Figure S2

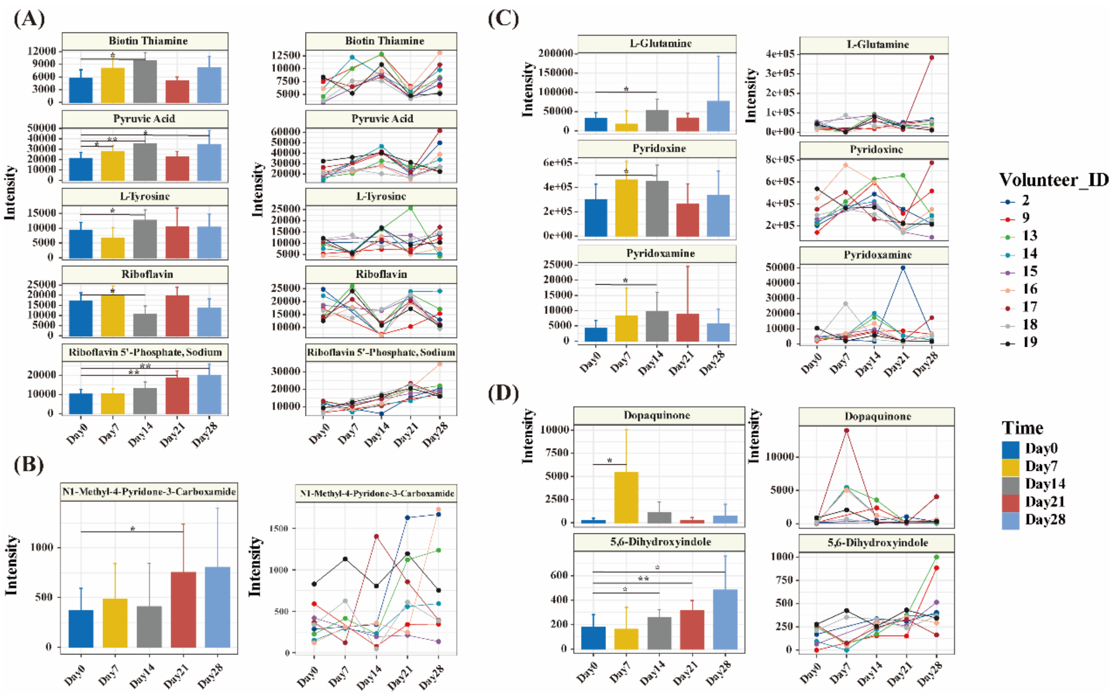


Changes in abundance of metabolites associated with (A) vitamin B1, B2 metabolism, (B) nicotinamide metabolism, (C) vitamin B6 metabolism, and (D) tyrosine metabolism. Statistical significance levels of data relative to baseline changes were assessed using the Wilcoxon paired test. Displayed results are p values. Unmarked indicates non-significance, * indicates $p \leqslant 0.05$, ** indicates $p \leqslant 0.01$.

### Figure S3

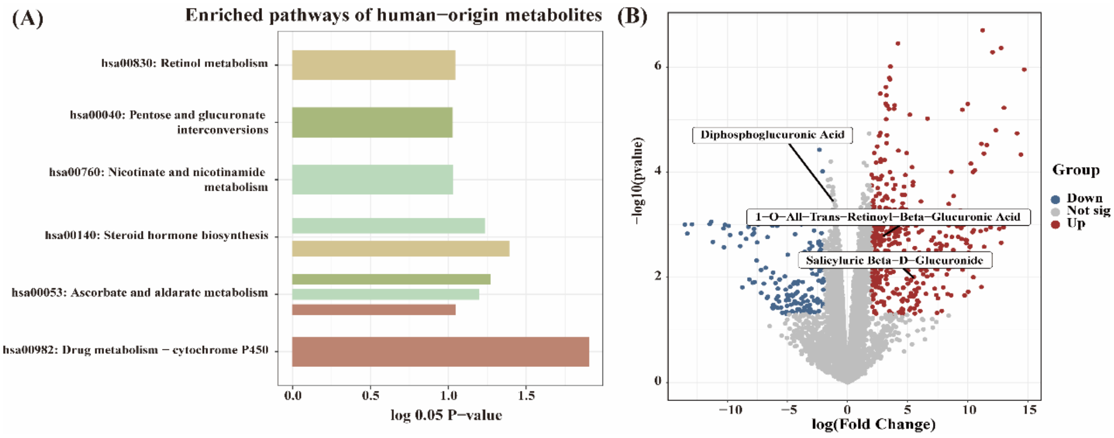


(A) enriched pathways of human-origin metabolites compared to baseline. (B) Volcano plot of differential metabolites relative to baseline on Day 7. The Limma package in R was utilized for identifying differential metabolites, employing paired tests and Benjamini-Hochberg correction for p-values. $\log FC \geqslant 2$ & adj.P $\leqslant 0.05$ was identified as “up regulated (up)” metabolites; $\log FC \leqslant -2$ & adj.P $\leqslant 0.05$ was identified as “down regulated (down)” metabolites; others were identified as “not significantly changed (not sig)” metabolites.
